# Initiation of Retina Regeneration By a Conserved Mechanism of Adult Neurogenesis

**DOI:** 10.1101/057893

**Authors:** Mahesh Rao, Dominic Didiano, James G. Patton

**Affiliations:** Department of Biological Sciences, Vanderbilt University, Nashville, TN

## Abstract

Retina damage or disease in humans often leads to reactive gliosis, preventing the formation of new cells and resulting in visual impairment or blindness. Current efforts to repair damaged retinas are inefficient and not capable of fully restoring vision. Conversely, the zebrafish retina is capable of spontaneous regeneration upon damage, using Müller glia (MG) derived progenitors. Understanding how zebrafish MG initiate regeneration may help develop new treatments that prompt mammalian retinas to regenerate. Here we show that inhibition of GABA signaling facilitates initiation of MG proliferation. GABA levels decrease following damage, and MG are positioned to detect the decrease. Using pharmacological and genetic approaches we demonstrate that GABAA receptor inhibition stimulates regeneration in undamaged retinas while activation inhibits regeneration in damaged retinas. GABA induced proliferation causes upregulation of regeneration associated genes. This is the first evidence that neurotransmitters control retina regeneration in zebrafish through an evolutionarily conserved mechanism of neurogenesis.

## Introduction

Retina damage or diseases, such as retinitis pigmentosa, result in a loss of retina cells and most often lead to blindness. A current effort to mitigate these effects involves intravitreal injections of stem cells or retina precursors, hoping for successful integration and connection to existing neuronal circuits (Barber et al., 2013; Hanus et al., 2016; MacLaren et al., 2006; Pearson et al., 2012; Santos-Ferreira et al., 2015). Though improving, these therapies, are very inefficient and not yet capable of restoring vision (Barber et al., 2013; Bringmann et al., 2006; Pearson, 2014; Pearson et al., 2010). An alternative method would be to prompt the retina to endogenously regenerate and replace lost cells.

Mammalian retinas do not possess the ability to regenerate following disease or damage. Instead, damage commonly results in reactive gliosis, preventing regeneration (Bringmann et al., 2006; Pearson, 2014). Zebrafish, however, mount a robust spontaneous regeneration response upon damage (Goldman, 2014). When damaged, Müller glia (MG) dedifferentiate, divide asymmetrically, and produce progenitor cells that are capable of restoring all lost cell types (Bernardos et al., 2007; Fausett and Goldman, 2006; Nagashima et al., 2013; Rajaram et al., 2014a; Rajaram et al., 2014b; Ramachandran et al., 2012; Thummel et al., 2008; Vihtelic and Hyde, 2000; Wan et al., 2012; Zhao et al., 2014). A number of key regulators that transition the retina through the stages of regeneration have been determined, but very little is known about what initiates the regeneration process. Because overall retina architecture and cells are largely conserved between fish and mammals, understanding how zebrafish initiate retina regeneration will help develop novel treatments or therapeutic targets for retina damage or diseases, especially treatments that target or induce regeneration from mammalian Müller glia.

Select regions of the mammalian central nervous system (CNS) are capable of adult neurogenesis, particularly the subgranular zone (SGZ) of the mouse hippocampus. Recently, the inhibitory neurotransmitter γ-aminobutyric acid (GABA) was shown to play an important role in regulating the proliferation of radial glia-like stem cells (RGLs) in the mouse hippocampus (Song et al., 2012). Synaptic input from glutamatergic granule cells regulates activity of GABAergic interneurons in the SGZ. When input from granule cells is low, decreases in extracellular GABA are detected by GABA_A_ receptors on RGLs, resulting in elevated proliferation. We sought to test whether this could be a mechanism that regulates regeneration in the damaged zebrafish retina. In the retina, photoreceptors (PRs) release glutamate onto GABAergic Horizontal cells (HCs). When PRs die they no longer stimulate HCs to release GABA. We hypothesize that MG detect the decrease in GABA and initiate regeneration in a response similar to activation of glial-like stem cells in the mouse hippocampus. We show here that disrupting GABA signaling causes spontaneous proliferation in undamaged zebrafish retinas and that increasing GABA signaling in damaged retinas suppresses regeneration. Our results indicate that GABA signaling is directly detected by MG, supporting an evolutionarily conserved mechanism of adult neurogenesis.

## Results

For MG to detect GABA released from HCs, their processes must be in close proximity. To test this, cross sections of retinas from two lines of zebrafish that express GFP in either MG (*Tg(gfap:GFP)^mi2001^*) or HCs (*Tg(lhx1a:EGFPf^303^*), were immunostained with antibodies against GABA, glutamic acid decarboxylase 65/67 (GAD65/67), or glutamine synthetase (GS). We found that HC and MG processes co-localize in the inner nuclear layer (INL) (Fig. 1 a-i). Closer examination of the INL in flat mounted *Tg(lhx1a:EGFP)^pt303^* retinas stained with GS showed that MG processes appear to wrap around HCs (Supplementary Fig. 1). Additionally, the subunit of the GABA_A_ receptor (GABRG2) that is essential for neurogenesis in the SGZ (Song et al., 2012) was detected on MG processes (Fig. 1j-m). This was validated by RT/PCR of sorted MG (data not shown) and by previous findings (Ramachandran et al., 2012).

**Figure 1.**
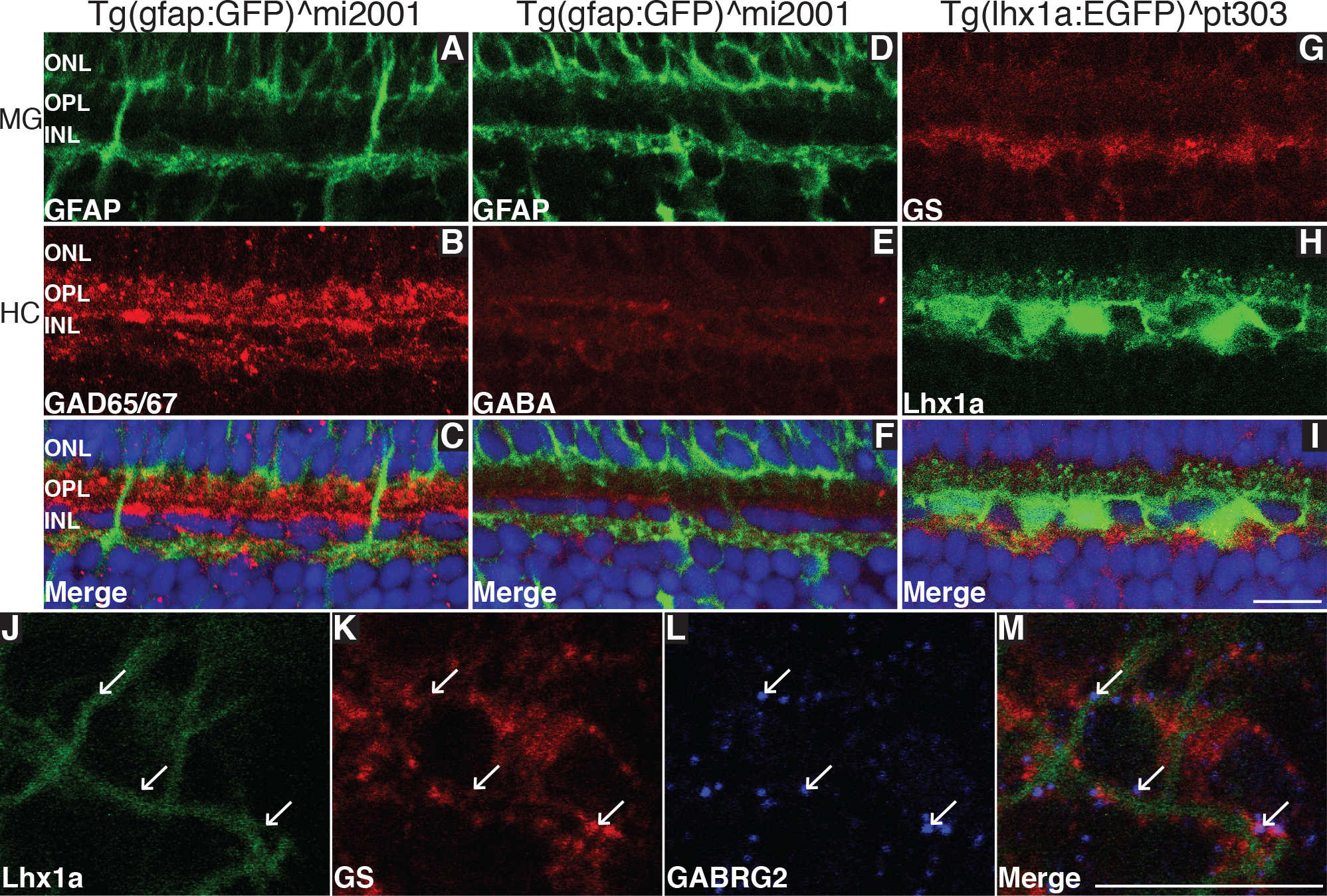
Close association of MG and HC processes in the INL.

*Tg(gfap:GFP)^mi2001^* and *Tg(lhx1a:EGFP)^pt303^* retina sections were stained for GAD 65/67 (a-C), GABA (D-F), or GS (G-I). Co-localization of MG and HC markers was observed in the INL. *Tg(lhx1a:EGFP)^pt303^*retinas were removed, stained for GS and GABRG2, and the area of co-localization imaged in flat mount (J-M). Arrows indicate GABRG2 puncta. Scale bar is 100μm.

To determine if GABA levels decrease after damage, *Tg(zop:nfsb-EGFP)^nt19^* fish were treated with metronidazole (MTZ) to destroy rod photoreceptors and retinas were harvested at different times of recovery (i.e. regeneration). It was found that MG begin to proliferate 52 hours after the end of MTZ treatment (Fig. 2).

**Figure 2.**
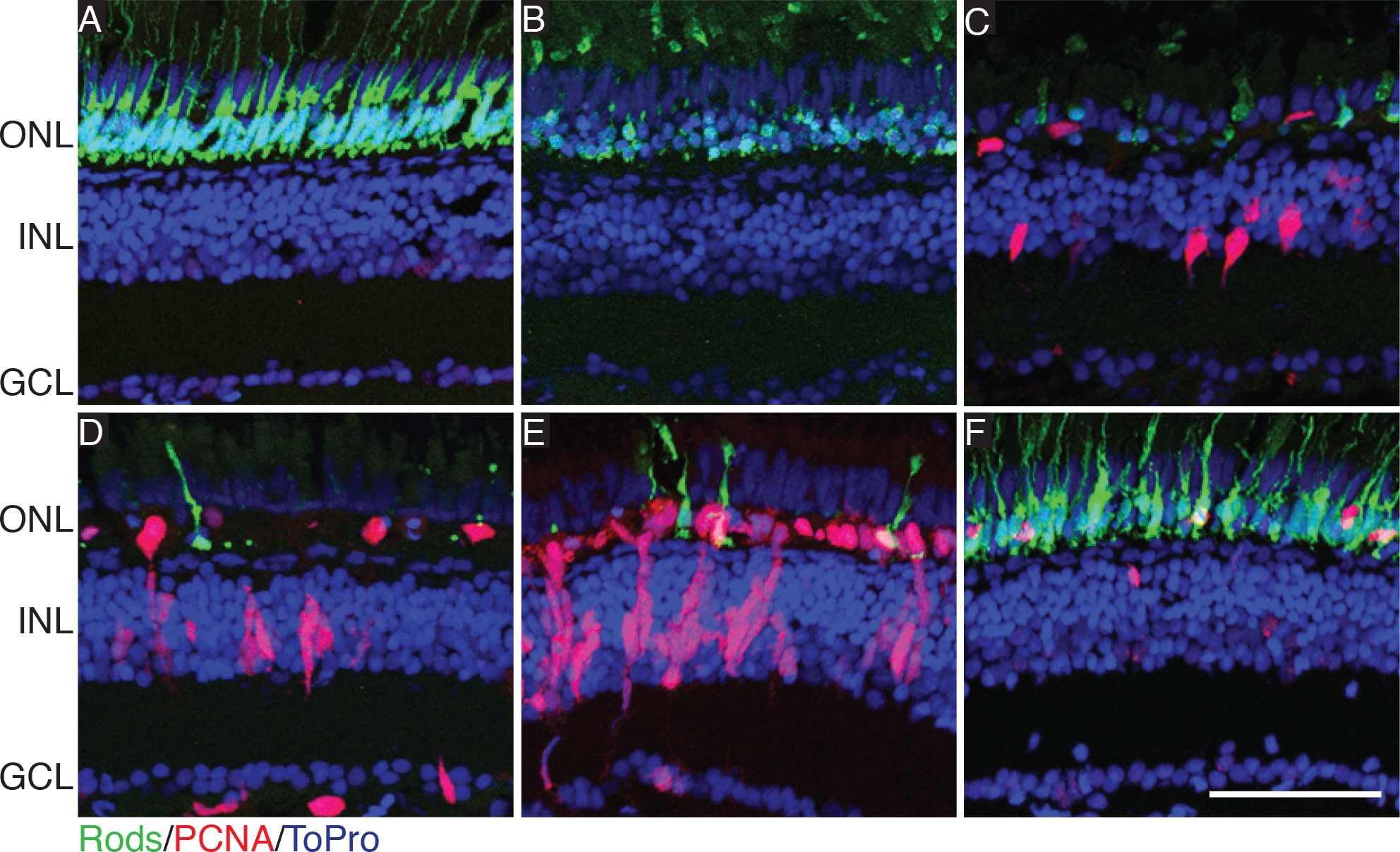
Timeline of regeneration in Tg(zop:nfsb-EGFP)^nt19^ after MTZ treatment.

*Tg(zop:nfsb-EGFP)^nt19^* fish were placed in egg water containing 10mM Metronidazole and treated for 24 hours, then returned to normal egg water to recover. Eyes were removed and proliferation assessed by PCNA staining. Times of recovery observed were pretreatment (A), 0h. recovery (B), 52h. recovery (C), 72h. recovery, 96h. recovery (D), and 28 days recovery (E). Scale bar is 100μm.

Whole retinas were then subjected to HPLC and the levels of GABA were measured. GABA levels were also significantly reduced 52 hours after MTZ treatment (Fig. 3). Together, these results support the idea that MG are poised to detect the decrease in GABA released from HCs that occurs after damage, causing them to regenerate. These findings prompted us to directly test whether altered GABA signaling initiates retina regeneration.

**Figure 3.**
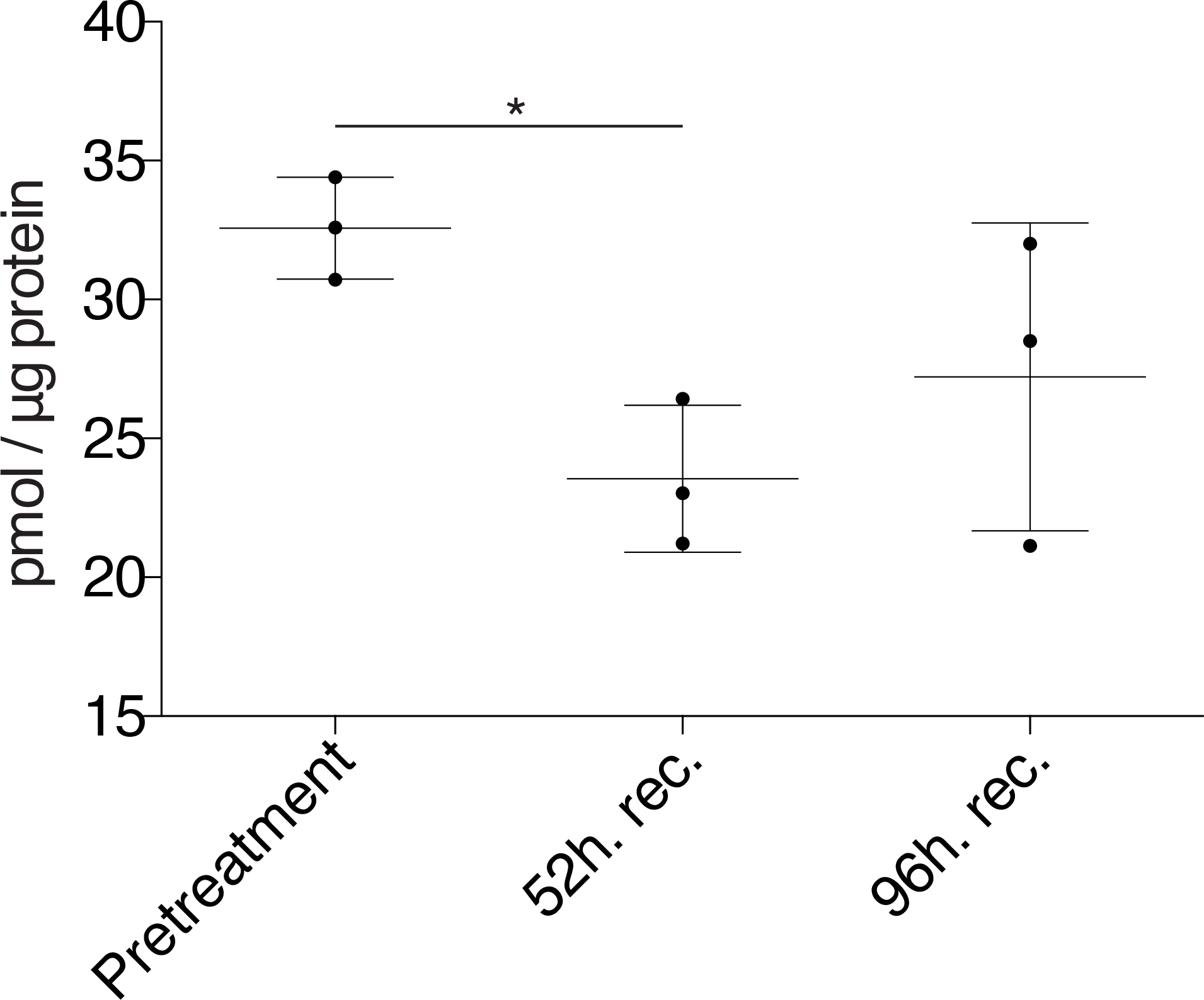
Whole retina GABA levels decrease following rod ablation.

*Tg(zop:nfsb-EGFP)^nt19^* fish were treated with MTZ and allowed to recover. Whole retinas were removed at indicated time points and levels of GABA were measured from lysates by HPLC. Levels of GABA were measured. A one-way ANOVA was used; Error bars = SD; * = p<0.05.

As a first test of the model, we injected a GABA_A_ antagonist (gabazine), into undamaged retinas and determined whether inhibition of GABA signaling would cause spontaneous proliferation (Fig. 4A). As early as 48 hours post injection (hpi) the number of proliferating cells, as detected by proliferating cell nuclear antigen (PCNA) expression, was significantly greater than both uninjected and PBS injected controls (Fig. 4B-D, Supplementary Fig. 2). The increase in proliferation was not a result of an increase in apoptosis (Supplementary Fig. 3). Proliferating cells were found in the INL and co-labeled with GS, indicating that MG were proliferating. Proliferating cells were found in clusters, which indicates a robust regenerative response with multiple divisions of MG-derived progenitor cells. The increase in spontaneous proliferation was dose dependent (Fig. 5) suggesting a specific effect via the GABA_A_ receptor.

**Figure 4.**
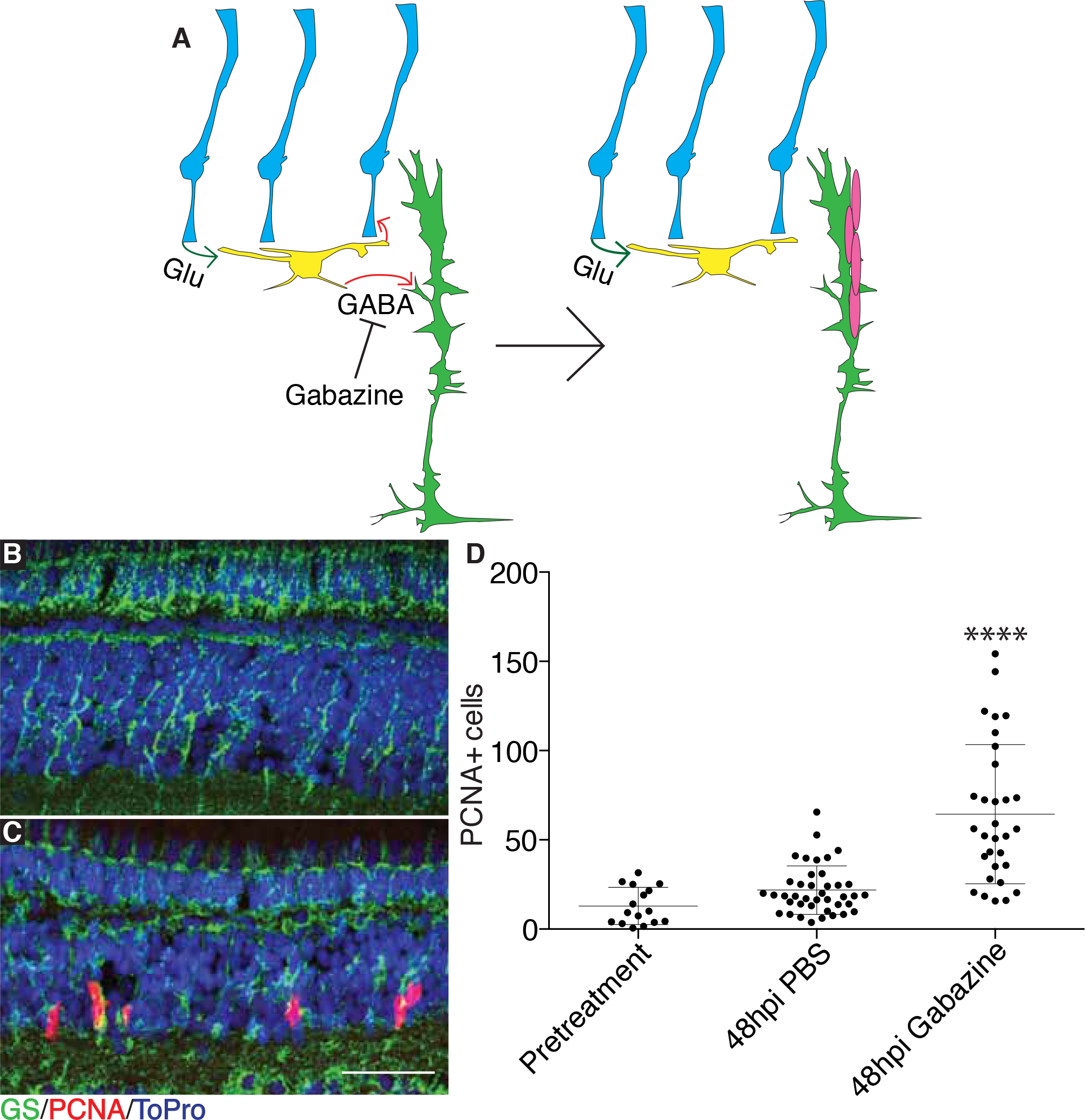
Gabazine injections cause time dependent spontaneous proliferation in undamaged retinas.

Model illustrating effects of gabazine injection on MG proliferation (A). WT eyes were injected with PBS (B) or 12.5 nmol gabazine (C) into one eye. Fish recovered for 48 hours after gabazine injections (B, C) before proliferation was measured. Representative images are small portions of entire retina. Proliferating cells were counted across whole sections by PCNA staining (D). Scale bar is 100μ. A one-way ANOVA was used; Error bars = SD;**** = p<0.0001.

**Figure 5.**
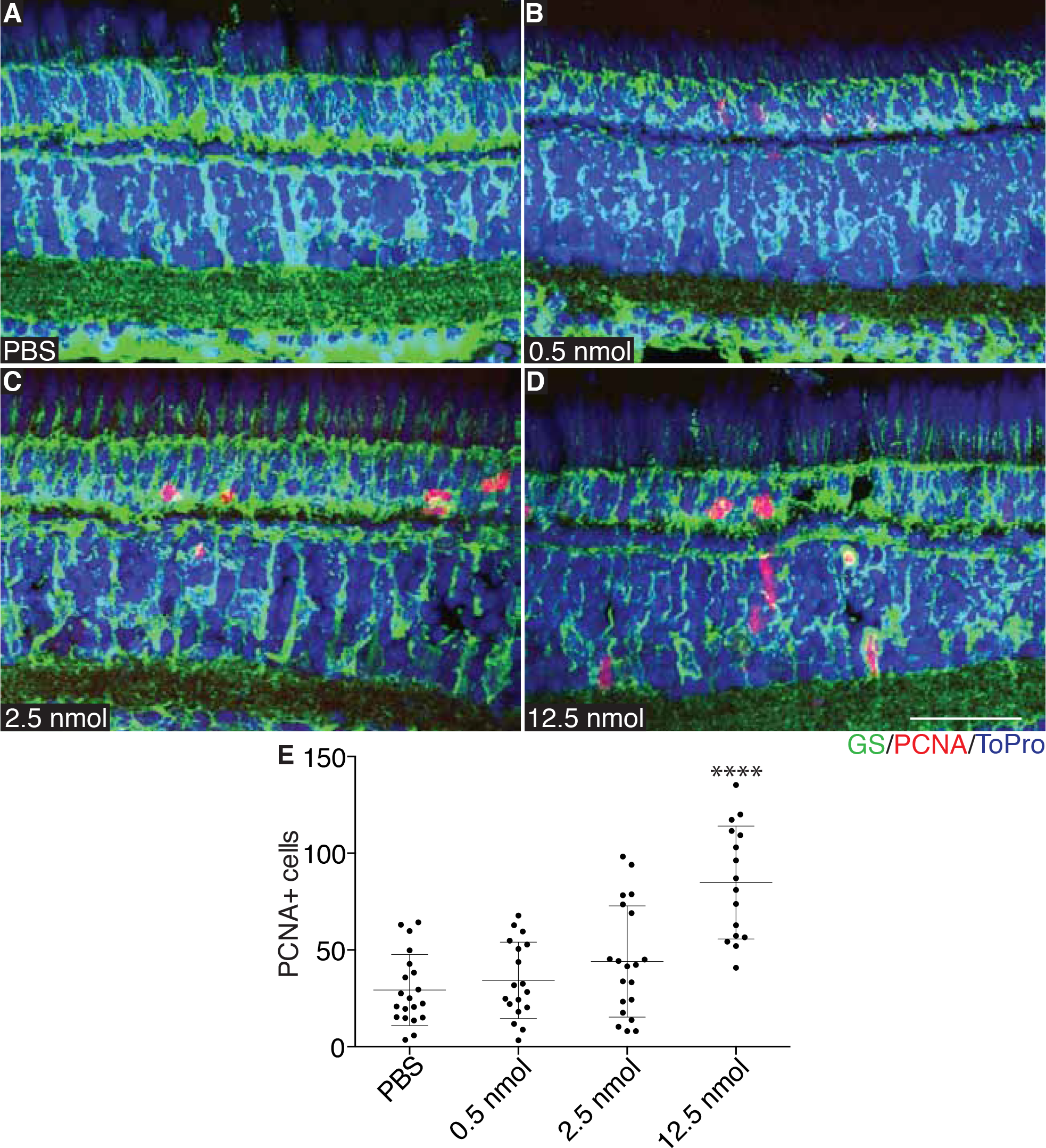
Gabazine induced spontaneous proliferation is dose dependent.

WT fish were injected with PBS (A), 0.5 nmol (B), 2.5 nmol (C), or 12.5 nmol (D) and proliferation measured at 48hpi by PCNA staining. Representative images are small portions of entire retina. Proliferating cells were counted across whole sections by PCNA staining (E). Representative images are small portions of total retina sections. Scale bar is 100pm. A one-way ANOVA was used; Error bars = SD; **** = p<0.0001.

Upstream of GABA signaling, a second prediction of the model is that inhibiting glutamate signaling should produce similar effects as inhibiting GABA (Fig. 6A). To test this, the AMPA receptor antagonist NBQX was injected into undamaged eyes and proliferation measured by PCNA staining. Proliferation was significantly greater at 48hpi but reached a maximum at 72hpi (Fig. 6B-D, Supplementary Fig. 4). As with gabazine injections, proliferation was not the result of increased apoptosis (Supplementary Fig. 5). Clusters of proliferating cells in the INL that co-labeled with GS were observed at 72hpi, indicating a robust regenerative response with multiple divisions of MG-derived progenitor cells. Furthermore, a dose dependent decrease in proliferation was also observed with NBQX injections (Fig. 7).

**Figure 6.**
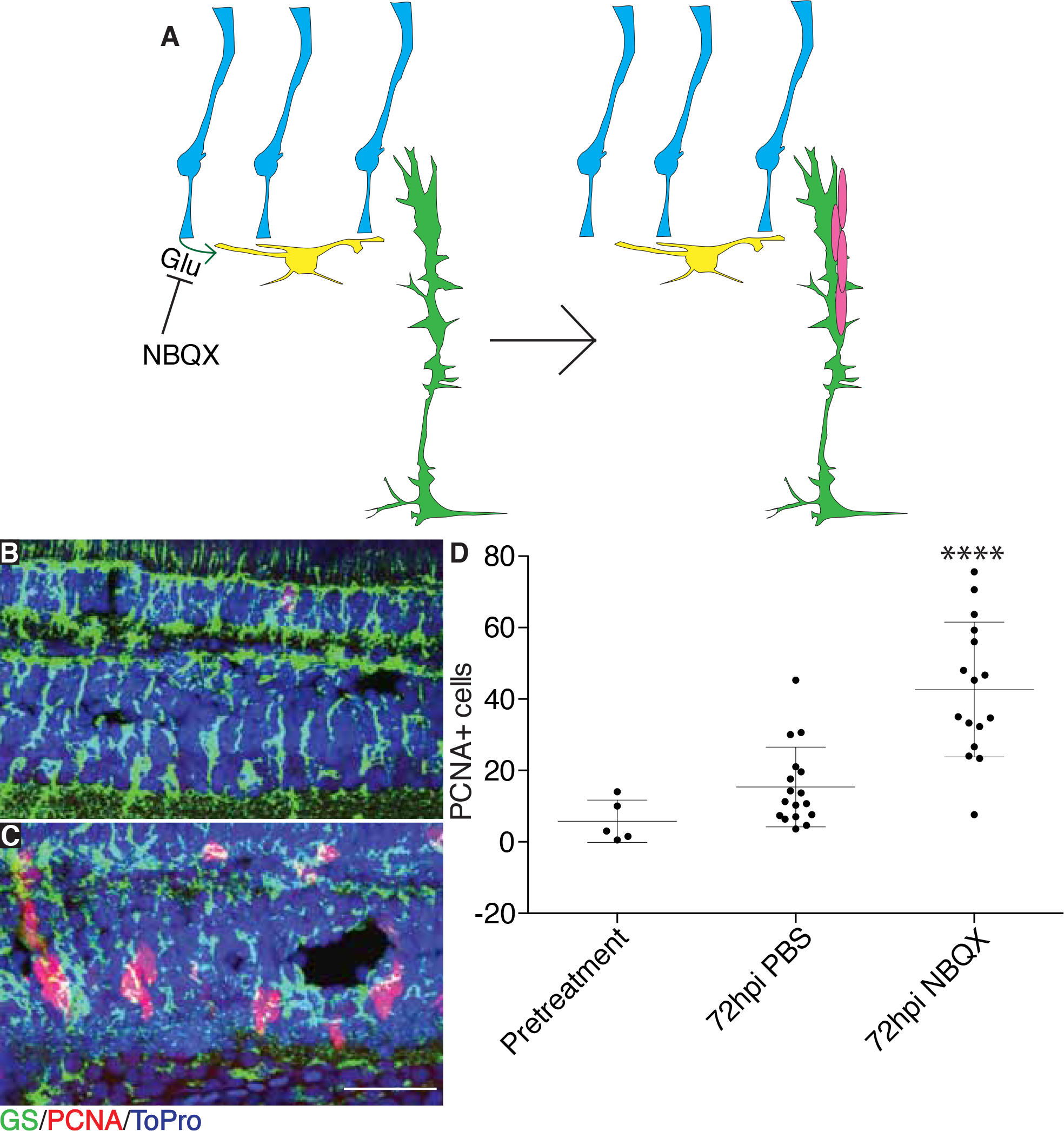
NBQX injections cause time dependent spontaneous proliferation in undamaged retinas.

Model illustrating effects of NBQX injections on MG proliferation (A). WT eyes were injected with PBS (C) or 25 nmol NBQX (C) into one eye. Fish recovered for 72 hours after NBQX injections (B, C) before proliferation was measured. Representative images are small portions of entire retina. Proliferating cells were counted across whole sections by PCNA staining (D). Scale bar is 100µm. A one-way ANOVA was used; Error bars = SD; ** = pp<0.01, **** = p<0.0001.

**Figure 7.**
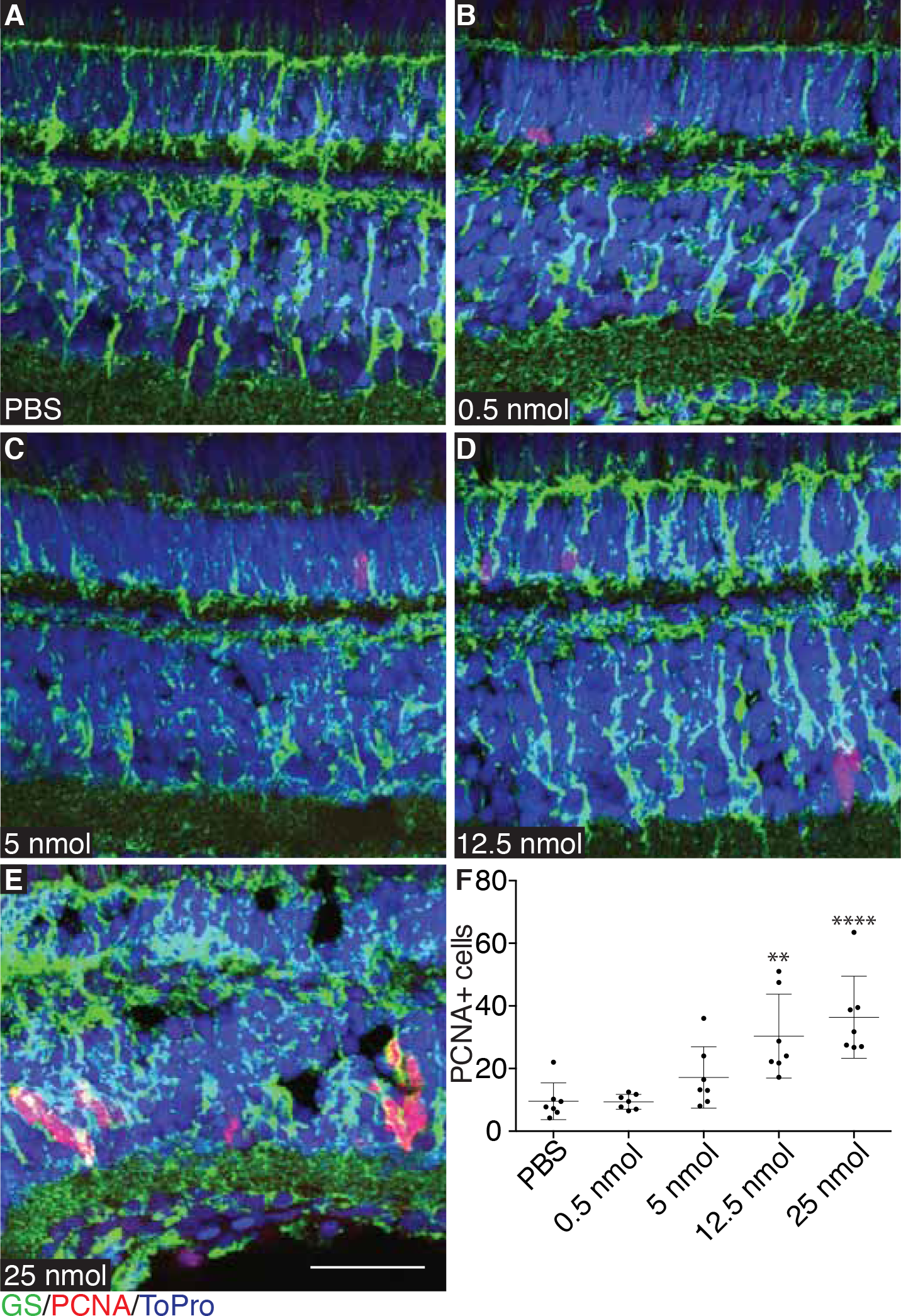
NBQX induced spontaneous proliferation is dose dependent.

WTfish were injected with PBS (A), 0.5 nmol (B), 5 nmol (C), 12.5 nmol (D), or 25 nmol (E) and proliferation measured at 72hpi by PCNA staining. Representative images are small portions of total retina sections. Proliferating cells were counted across whole sections by PCNA staining (F). Representative images are small portions of total retina sections. Scale bar is 100µm. A one-way ANOVA was used; Error bars = SD; ** = p<0.01, **** = p<0.0001.

In order to verify that proliferation resulting from gabazine or NBQX injections accurately replicates regeneration, markers of regeneration were analyzed following injections. Drugs were injected into *Tg(tuba1a:GFP)*, and *Tg(her4:dRFP)* fish, marking activation of α-tubulin 1a and Notch signaling, respectively. Both genes have been shown to be upregulated during regeneration (Fausett and Goldman, 2006; Hayes et al., 2007; Ramachandran et al., 2010;Wan et al., 2012). After injection, both were found to be upregulated and associated with PCNA expressing cells (Fig. 8) suggesting that drug induced proliferation is accurately replicating what is observed during damaged-induced retina regeneration.

**Figure 8.**
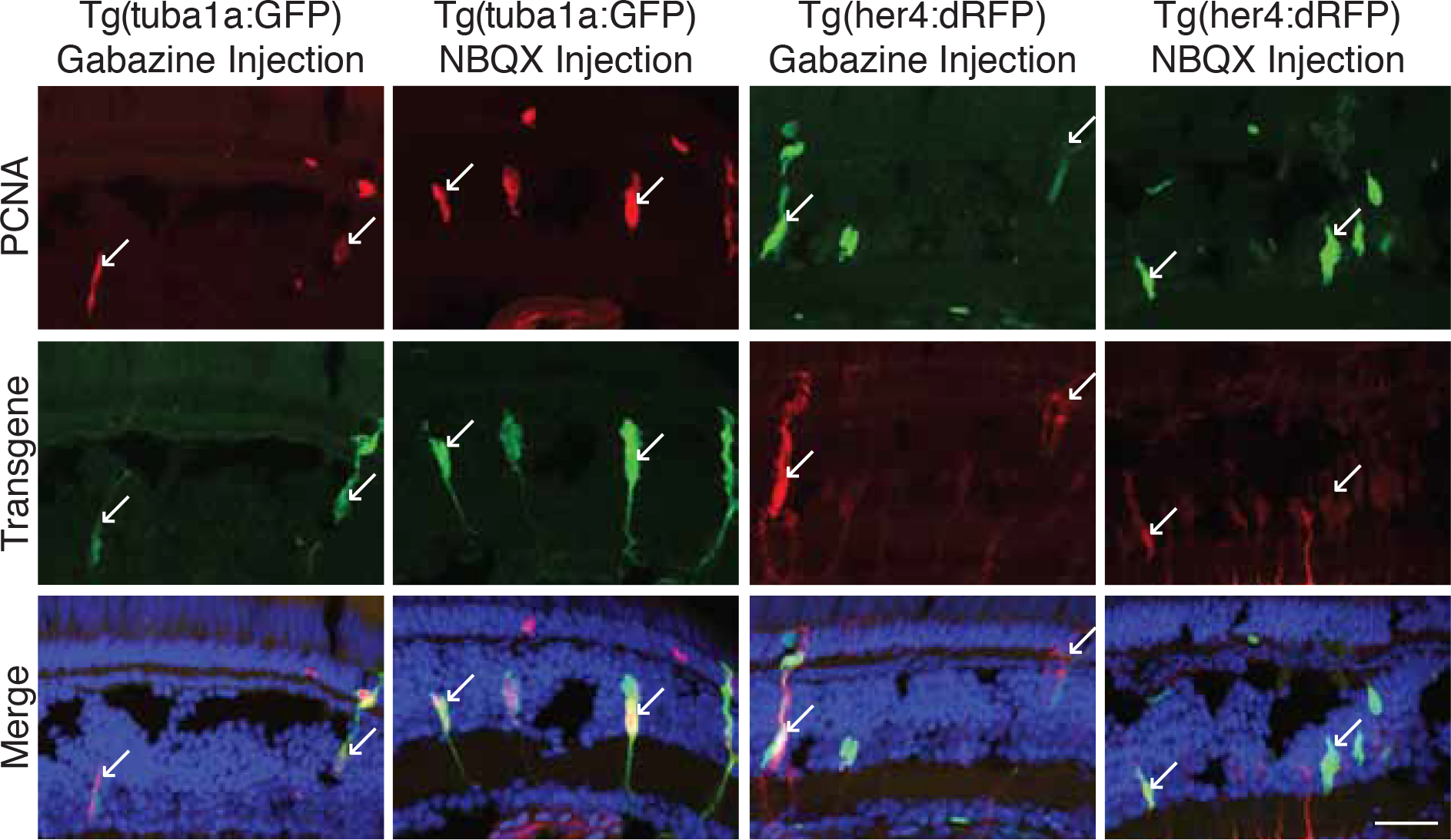
Injection of gabazine or NBQX into undamaged eyes causes upregulation of factors associated with regeneration.

Gabazine or NBQX was injected into one eye of *Tg(tuba1a:GFP)* or *Tg(her4:dRFP)* fish. Fish were allowed to recover for 72 hours, after which retinas were removed and stained for PCNA. Both GFP and dRFP expression colabeled with PCNA. Arrows indicate colocalization of transgene and PCNA. Scale bar is 100µm.

The above experiments showed an increase in proliferation without damage. A converse set of experiments was devised to determine whether activating the GABA_A_ receptor or the AMPA receptor would suppress regeneration after damage (Fig. 9A). To test this, *Tg(zop:nfsb-EGFP)^nt19^* fish were damaged with MTZ and eyes injected with either muscimol, a GABA_A_ receptor agonist, or AMPA, a glutamate receptor agonist. Injected retinas were collected at 52 hours after MTZ treatment, when MG proliferation begins (Fig. 2). Both injections showed a significant decrease in proliferating cells compared to PBS injections, as measured by PCNA expression (Fig. 9B-F).

**Figure 9.**
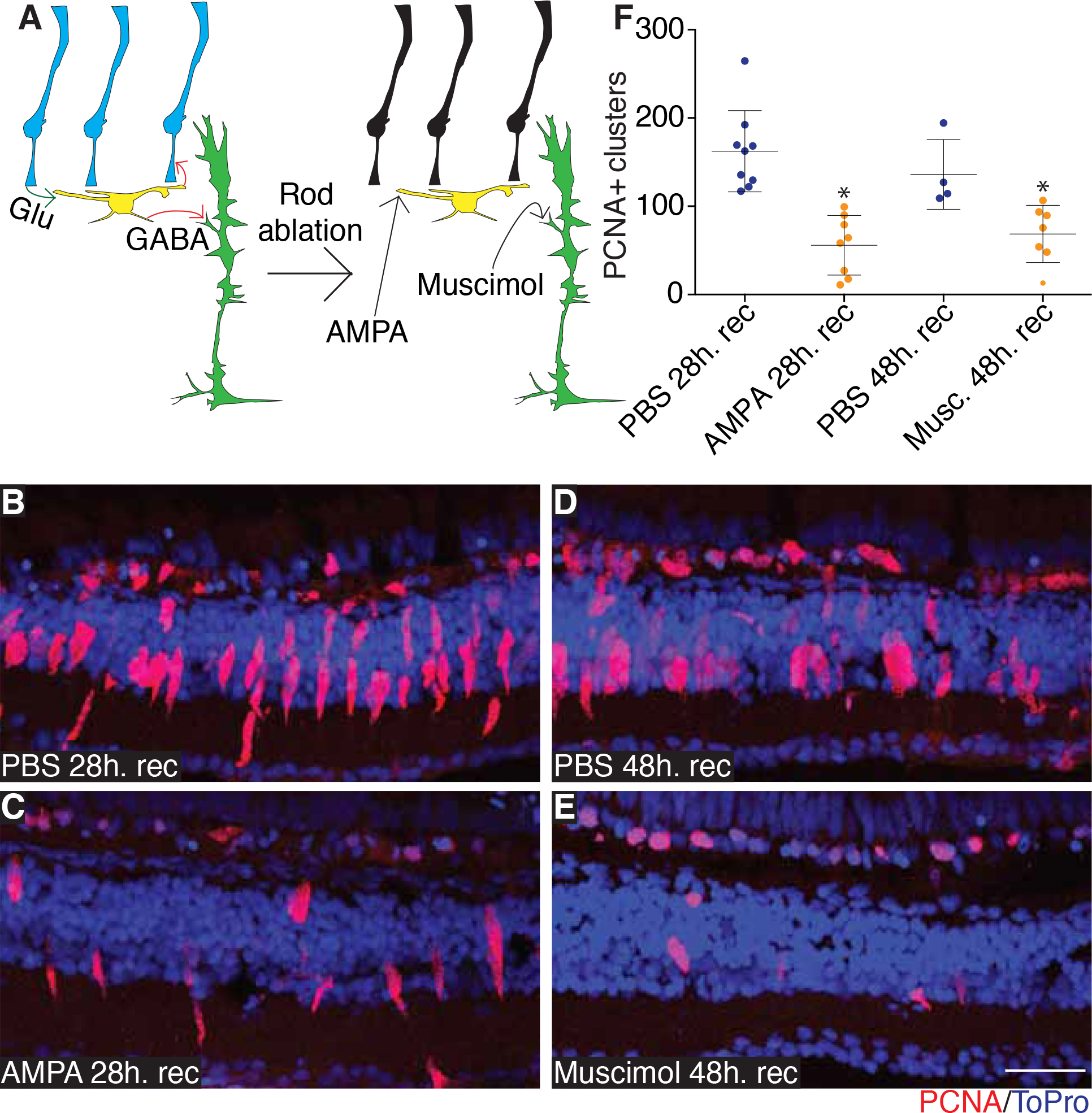
AMPA and Muscimol injections suppress regeneration in damaged retinas.

Model illustrating effects of muscimol and AMPA injections on MG proliferation (A). *Tg(zop:nfsb-EGFP)^nt19^* fish were treated with 10mM metronidazole for 24 hours, then allowed to recover. Fish were then anesthetized and injected with either AMPA/PBS control at 28h. recovery or muscimol/PBS control at 48h. recovery. Injected eyes were removed at 52h. recovery. Proliferation was assessed by PCNA staining (B-E). Representative images are small portions of the entire retina. Clusters of proliferating cells were measured across entire sections (F). Scale bar is 100μm. A Students t-test was used; Error bars = SD; * = p<0.05.

One caveat to the above experiments is that the various agonists and antagonists could be acting indirectly to cause MG proliferation. To address this, we used a genetic approach by creating a construct expressing a dominant negative version of the zebrafish γ^2^ subunit of the GABA_A_ receptor (DNγ2) following an identical human mutation that underlies an inherited form of epilepsy (Harkin et al., 2002; Kang et al., 2009). The zebrafish glial fibrillary acidic protein (GFAP) promoter was used to drive expression of an mCherry tagged version of the DNγ2 isoform in MG. Injection and electroporation of this construct into *Tg(gfap:GFP)^mi2001^* fish was performed followed by analysis of proliferating cells that co-localize with mCherry, GFP, and PCNA. There was a significant increase in the total number of proliferating cells, marked by PCNA and co-localizing with GFP and mCherry (Fig. 10, Supplemental Fig. 6). This indicates that MG directly respond to GABA.

**Figure 10.**
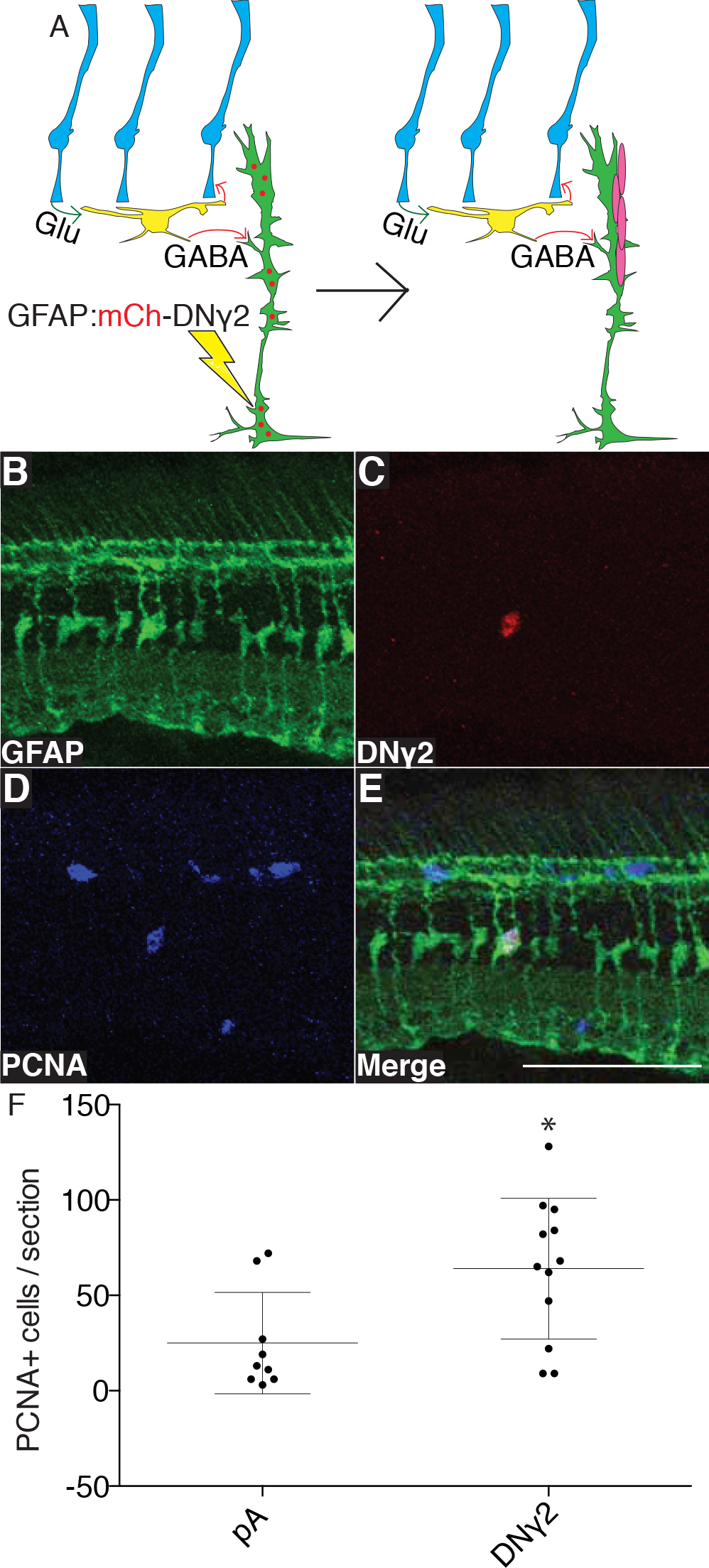
Expression of DNγ2 in MG of undamaged retina causes increased proliferation.

Model illustrating effects of electroporation of DNγ2 into MG on proliferation (A). A GFAP:mCh-DNγ2 construct was electroporated into one retina of undamaged *Tg(gfap:GFP)^mi2001^* fish. GFP expression (B), mCh expression (C), and staining for PCNA (D) all co-labeled in the same cell (E). Total number of PCNA expressing cells was measured (F). Scale bar is 100nm. A Students t-test was used; Error bars = SD; * = p<0.05.

## Discussion

Our data support a novel mechanism in which decreased GABA is sensed by MG to initiate retina regeneration. Photoreceptor death results in a decrease in GABA, and MG are poised to detect this decrease by their close association with HCs and their expression of GABA_A_ receptors. These findings indicate that regulation of neurogenesis in the mouse hippocampus is a conserved mechanism across structures and organisms and strongly suggests this is a basic archetype for eliciting neurogenesis. GABA has been found to be involved in neurogenesis in other areas of the CNS, specifically the SGZ and subventricular zone (SVZ). Multiple reports have indicated an importance for GABA in proliferation of progenitor cells (Braun and Jessberger, 2014; Giachino et al., 2014; Liu et al., 2005; Pallotto and Deprez, 2014; Ramirez et al., 2012; Song et al., 2013; Tozuka et al., 2005). This may also be true in the retina, where elevated GABA signaling may promote progenitor cell proliferation or differentiation but suppress MG proliferation, providing directionality for the regeneration process.

Our data suggest that PR damage is communicated via GABA, based on the timing of events in different experiments. Nevertheless, the possibility exists that MG directly sense changes in glutamate as well. Maximum proliferation for gabazine injections was observed at 48hpi while maximum proliferation for NBQX injections was observed at 72hpi. This suggests that GABA affects MG more proximally than glutamate. Furthermore, muscimol injection into damaged retinas only produced an effect when injected at 48 hours after MTZ treatment, while AMPA injections only produced an effect when injected at 28 hours after MTZ treatment. Injecting muscimol earlier or AMPA at later times did not cause significant changes in proliferation (data not shown). The overall timing best supports the idea that GABA acts directly on MG and glutamate is upstream.

Previous studies have suggested that TNFα (Nelson et al., 2013), Notch(Conner et al., 2014), leptin, and interleukin 6 (IL-6)(Zhao et al., 2014), are involved in initiating retina regeneration in zebrafish. It is possible that these and other, as yet undiscovered, signals act synergistically to mount a full, robust regenerative response. However, even though TNFα, Notch, leptin, and IL-6 are all relatively early markers of regeneration, it is not clear what signals induce their expression, especially because numerous gene expression changes accompany differential expression of these factors. An attractive hypothesis based on our data is that decreased GABA is the primary signal for retina regeneration initiation and other signals follow to act synergistically. This is supported by the fact that injection of gabazine or NBQX causes an upregulation of the Notch reporter Her4, suggesting that GABA is upstream of Notch signaling. This is also in line with earlier studies that showed that Notch signaling is important for later stages of regeneration and development (Hayes et al., 2007; Karl et al., 2008; Kassen et al., 2007; Olena et al., 2015; Raymond et al., 2006). In addition, leptin mRNA was observed to increase following injury, suggesting that it is induced by some other signal. Interestingly, Il-6 mRNA was not detected in regenerating retinas, indicating that its source originates from outside the retina and may be prompted to increase only after damage occurs (Zhao et al., 2014). Inflammatory signals like TNFα and Il-6 may be released by endogenous immune cells (e.g. microglia(Fischer et al., 2014)) or those invading from the vasculature, after damage.

Retinitis pigmentosa and age related macular degeneration arise from dysfunction and death of photoreceptors. We have focused on photoreceptor regeneration but the questions remains how bipolar, amacrine, or ganglion cells regenerate. It may be that feedback mechanisms are in place where HC activity would be affected by the death of other cells. For example, dopaminergic amacrine cells have been found to have processes that project to the HC layer and may also affect HC activity (Herrmann et al., 2011). There are likely other mechanisms by which the retina senses bipolar, amacrine, or ganglion cell death as well. MG processes appear to surround the cell bodies of amacrine and ganglion cells (data not shown), suggesting a different mechanism to sense cell death, such as juxtacrine or paracrine signaling. These mechanisms may also be involved in sensation of PR and HC death. Identifying a method to induce spontaneous MG proliferation and the production of progenitor cells may be sufficient if the new progenitors can then follow endogenous cues to differentiate into whatever cell is needed. New therapies that activate MG in this manner could lead to robust endogenous regeneration and counteract many retina diseases. These might include both agonists and antagonists of neurotransmitter signaling but could also be targeted at intracellular cascades downstream of GABA binding as well as other factors (Jagasia et al., 2009; Quadrato et al., 2012; Quadrato et al., 2014; Ramirez et al., 2012) (Andang et al., 2008; Fernando et al., 2011)

Lastly, the question remains why teleost retinas have maintained a robust regenerative response while mammalian retinas are largely incapable of repair. The cellular organization found in zebrafish may be a key difference. In zebrafish, the HCs form a monolayer that is separated from the rest of the INL by the network of HC and MG processes, observed in the current study (Fig. 1). A consequence of this organization is that the HCs contain processes that project into the INL as well as into the outer plexiform layer (OPL). In mice, however, the HCs are comingled with other cells in the INL and only contain projections into the OPL(Matsuoka et al., 2012; Poche et al., 2007). Investigating HC development and potential interactions between HCs and MG could greatly inform about how MG respond to damage in the mammalian retina. It may be that MG are inefficient or blocked from detecting changes in GABA after damage but perhaps alteration of signaling by pharmacological agents such as gabazine could push MG down a regenerative path.

## Acknowledgments

We wish to thank members of the Patton lab, Elizabeth Beilharz, and Alissa Guarnaccia for help and advice. Transgenic zebrafish lines were shared by Pamela Raymond (*Tg(gfap:GFP)^mi2001^*), David Hyde (*Tg(zop:nfsb-EGFP)^nt19^),* Neil Hukriede (*Tg(lhx1a:EGFPf^303^),* Daniel Goldman (*Tg(tuba1a:GFP)*), and Ajay Chitnis (*Tg(her4:dRFP)*). This work was supported by grants from the National Institutes of Health RO1 EY024354 and R21 EY019759 to JGP with additional support from the Stevenson and Gisela Mosig endowments to Vanderbilt University.

## Author Contributions

MBR, DD, and JGP conceived, designed, and performed all experiments and wrote the paper.

## Competing Financial Interests

The authors declare no competing financial interests.

## Materials and Methods

### Zebrafish lines and Maintenance

Zebrafish lines used in this study include *Tg(gfap:GFP)^mi2001^* (Bernardos and Raymond, 2006), which marks differentiated MG, *Tg(zop:nfsb-EGFP)^nt19^* (Montgomery et al., 2010), which marks rods and is used for rod ablation, *Tg(lhx1a:EGFP)^pt303^* (Swanhart et al., 2010), which marks HCs, *Tg(tuba1a:GFP)* (Fausett and Goldman, 2006), which marks dedifferentiated MG and progenitor cells, and *Tg(her4:dRFP)* (Yeo et al., 2007), which marks notch activated cells. All fish were maintained in a 14:10 light:dark (L:D) cycle at 28^°^C unless otherwise noted.

### Metronidazole induced rod damage

Rod ablation was induced similar to previously established protocols (Montgomery et al., 2010). Briefly, *Tg(zop:nfsb-EGFP)^nt19^* transgenic zebrafish were transferred to egg water containing 10mM metronidazole (MTZ) for 24 hours in darkness at 28^°^C. Fish were then transferred to normal egg water and returned a 14:10 L:D cycle for recovery. The extent of regeneration was assayed at the indicated times post recovery after MTZ treatment.

### HPLC analysis of GABA

Whole retinas were dissected from *Tg(zop:nfsb-EGFP)^nt19^* transgenic zebrafish following recovery from MTZ treatment. Protein was extracted from 10 whole retinas for each time point and subjected to High-Performance Liquid Chromatography (HPLC). The levels of each amino acid and derivations were quantified.

### Drug injections

Different neurotoxins were injected into the vitreous using a protocol adapted from previous studies (Rajaram et al., 2014a; Rajaram et al., 2014b; Thummel et al., 2008). The drugs included gabazine (25mM; Sigma S106), nitro-2, 3-dioxobenzoquinoxaline-sulfonamide (NBQX) (50mM; Abcam ab210046), muscimol (5mM; Sigma M1523), and α-amino-3-hydroxy-5-methyl-4-isoxazolepropionic acid (AMPA) (10µM; Sigma A9111). Briefly, zebrafish were anesthetized in 0.016% tricaine, an incision was made in the sclera with a sapphire knife, and a blunt end 30 gauge needle inserted. 0.5µL were injected into one eye of adult zebrafish. Fish were immediately placed into a recovery tank, times indicated are hours of recovery.

### Immunohistochemistry and TUNEL labeling

Zebrafish were euthanized in 0.08% tricane and whole eyes were removed and fixed in 9:1 ethanolic formaldehyde (PCNA staining) or 4% paraformaldehyde (all other staining) overnight. Eyes were then washed in PBS and cryoprotected in 30% sucrose for 4 hours at room temperature. Eyes were then transferred to a solution containing 2 parts OCT and 1 part 30% sucrose overnight followed by transfer to 100% OCT for 2 hours and then embedded in OCT for cryosectioning. Antibodies used were PCNA (Sigma, P8825; Abcam, ab2426), Glutamine Synthetase (Millipore, mab302), GABA (Sigma, A0310), GABAa receptor gamma 2 subunit (Novus Biologicals, NB300-151), GAD65+GAD67 (Abcam, ab11070), GFP (Torrey Pines BioLabs, TP401), mCherry (Novus Biologicals, NBP1-96752). TUNEL labeling was performed following IHC. The In Situ Cell Death Detection Kit, TMR red (Roche Applied Sciences, 12156792910) was used to detect apoptotic cells.

### Design of Dominant Negative γ2 and electroporation

A dominant negative form of the γ2 subunit of the GABA_A_R was previously characterized in humans (Harkin et al., 2002; Kang et al., 2009). The mutation is in a conserved position in zebrafish and generates a premature codon. A plasmid containing the mRNA sequence of γ2 until the premature stop codon was created by GeneArt^®^. The sequence was cloned into a Tol2 backbone to create the vector GFAP:mCh(no-stop)-DNγ2. A control vector GFAP:mCh(no-stop)-pA was also created. Both constructs were electroporated into retinas following a protocol adapted from previous studies (Rajaram et al., 2014a; Rajaram et al., 2014b; Thummel et al., 2008). Briefly, fish were anesthetized, the outer cornea was removed, an incision was made in the sclera with a sapphire knife, and a blunt end 30 gauge needle was inserted into the vitreous. 0.5μL of plasmid DNA at a concentration of 2 ng/μL was injected into the vitreous of one eye. Anesthetized fish were allowed to recover, re-anesthetized, and then injected eyes were electroporated (50 V/pulse, 4 pulses, 1 second intervals between pulses). Treated fish were placed into recovery tanks for the times indicated.

### Statistical Analysis

A two-tailed Students t-test analysis was performed when comparing two means and a one-way ANOVA analysis was performed when comparing 3 or more means. The test used is indicated in each figure as well as the number of eyes measured. If possible, experiments contained a single clutch of fish that were divided among different treatment groups. When multiple clutches were required, fish were mixed, then distributed among treatment groups to reduce bias. 3 sections from one eye were measured and the resulting values averaged to arrive at the reported value.

